# Toxicity of JUUL Fluids and Aerosols Correlates Strongly with Nicotine and Some Flavor Chemical Concentrations

**DOI:** 10.1101/490607

**Authors:** Esther E. Omaiye, Kevin J. McWhirter, Wentai Luo, James F. Pankow, Prue Talbot

**Affiliations:** Environmental Toxicology Graduate Program, University of California Riverside, California, USA; Department of Civil and Environmental Engineering, Portland State University, Portland, Oregon, USA; Department of Chemistry Portland State University. Portland, Oregon, USA; Department of Molecular, Cell and Systems Biology, University of California, Riverside, California, USA

**Keywords:** Nicotine, JUUL, EC fluids, cytotoxicity, flavor chemicals, GC/MS

## Abstract

While JUUL electronic cigarettes (ECs) have captured the majority of the EC market with a large fraction of their sales going to adolescents, little is known about their cytotoxicity and potential effects on health. The purpose of this study was to determine flavor chemical and nicotine concentrations in the eight currently marketed pre-filled JUUL EC cartridges (“pods”) and to evaluate the cytotoxicity of the different variants (e.g., “Cool Mint” and “Crème Brulee”) using in vitro assays. Nicotine and flavor chemicals were analyzed using gas chromatography/mass spectrometry in pod fluid before and after vaping and in the corresponding aerosols. 59 flavor chemicals were identified in JUUL pod fluids, and three were >1 mg/mL. Duplicate pods were similar in flavor chemical composition and concentration. Nicotine concentrations (average 60.9 mg/mL) were significantly higher than any EC products we have analyzed previously. Transfer efficiency of individual flavor chemicals that were >1mg/mL and nicotine from the pod fluid into aerosols was generally 35 - 80%. All pod fluids were cytotoxic at a 1:10 dilution (10%) in the MTT and neutral red uptake assays when tested with BEAS-2B lung epithelial cells. Most aerosols were cytotoxic in these assays at concentrations >1%. The cytotoxicity of aerosols was highly correlated with nicotine and ethyl maltol concentrations and moderately to weakly correlated with total flavor chemical concentration and menthol concentration. Our study demonstrates that: (1) some JUUL flavor pods have high concentrations of flavor chemicals that may make them attractive to youth, and (2) the concentrations of nicotine and some flavor chemicals (e.g. ethyl maltol) are high enough to be cytotoxic in acute in vitro assays, emphasizing the need to determine if JUUL products will lead to adverse health effects with chronic use.

## INTRODUCTION

While cigarette smoking is declining in many countries, youth and adult use of e-cigarettes (ECs) has increased.^1–3^ and EC sales are estimated to reach 3.6 billion dollars in 2018.^4^ To appeal to consumers and improve nicotine delivery, ECs have evolved since their introduction into world markets about 10 years ago. Although original models looked similar to tobacco cigarettes and were often termed “cig-a-likes”,^5^ some highly evolved models have large tanks and batteries with features that allow power control.^6^

The JUUL brand is one of the newer entries into the EC market and is more similar to the “cig-a-like” products than to recently available tank/box mod styles.^7^ JUUL has spurred the development of many competing single pod style atomizers designed to be used with refill fluids containing nicotine salt.^8,9^ In June 2018, in the US, it was estimated that about 68% of current EC sales are JUUL products.^10^ Middle and high school students, as well as young adults, make up a large fraction of JUUL consumers.^11^ This demographic may be attracted to JUUL in part because of its appealing compact design, which resembles a USB drive, and its ability to create relatively small clouds of aerosol making its use indoors and in schools difficult to detect.^12^ Unlike many other EC fluids, JUUL products contain high concentrations of nicotine and sufficient acid to protonate most of the nicotine; lower free-base nicotine levels have been associated with increased palatability on inhalation.^13–15^

The JUUL utilizes pre-filled EC fluid “pods”, originally sold exclusively by JUUL, but now offered by third parties. JUUL currently sells eight flavors of pods, which can be characterized as minty (“Cool Mint” and “Classic Menthol”), fruity (“Mango”, “Fruit Medley” and “Cool Cucumber”), sweet (“Crème Brulee”), and tobacco (“Classic Tobacco” and “Virginia Tobacco”). In spite of their sudden surge in popularity, relatively little is known about the chemicals delivered by JUUL products. We have previously shown that many EC refill fluids contain very high concentrations of flavor chemicals^16,17^ and that these concentrations are cytotoxic when tested in vitro with lung cells.^17–20^

The purposes of this study were to: (1) quantify nicotine concentrations in the eight products offered by JUUL and compare these concentrations to other EC products, (2) identify and quantify the flavor chemicals in the eight flavor pods, (3) determine the transfer efficiency of nicotine and flavor chemicals into aerosols, and (4) test these products for cytotoxicity *in vitro* using human lung cells.

## MATERIALS AND METHODS

### Purchase of JUUL Products

The five original flavors of JUUL pods and three “limited edition” flavors were purchased online from the manufacturer’s USA website. These were “Cool Mint”, “Crème Brulee”, “Mango”, “Fruit Medley”, “Virginia Tobacco”, “Cool Cucumber”, “Classic Menthol” and “Classic Tobacco” (Supporting Information, S1). Products were inventoried and stored at room temperature until used. According to the manufacturer’s label information, each JUUL Pod flavor contains 0.7 mL of flavored fluid at 5% nicotine.

### Aerosol production and capture using an impinger method

Aerosol generated from pod fluids was bubbled through and captured in either isopropyl alcohol for flavor chemical and nicotine analysis or basal cell culture medium for cytotoxicity evaluation. These aerosols captured in a fluid will be referred to as “aerosol” in the remainder of the paper. Aerosols produced from different pod flavors were collected at room temperature in two tandem 125 mL impingers, each containing 25 mL of isopropanol or basal cell culture medium. A JUUL EC (battery and pre-filled pod) connected to a Cole-Parmer Masterflex L/S peristaltic pump was puffed using a 4.3 sec puff duration^21^ and an air flow rate of 10 – 13 mL/sec. To avoid dry puffing, only ¾ of the pod fluid was vaped. The pods were weighed before and after aerosol production to collect at least 15 mg/mL for GC/MS analysis. Aerosol solutions were stored at −20 °C until shipped to Portland State University for analysis.

For the MTT assay, 6 total puff equivalents or TPEs (1 TPE = 1 puff/milliliters of culture medium) aerosol solutions were prepared in BEAS-2B basal medium and supplements were added after aerosol production. The complete medium was passed through a 0.2 μ filter, and aliquots were stored at −80 °C until testing. Aerosols were tested at 0.02, 0.06, 0.2, 0.6, 2 and 6 TPE. To convert from TPE to percentage of the concentration of the pod fluid, the pod weight difference before and after aerosol collection was utilized to obtain the mg of fluid consumed. The weight (grams) of fluid consumed/puff of aerosol was calculated, and the density of the pod fluid was determined. Then the grams/puff were converted to milliliters using the density values. Finally, the percent for concentrations used in the aerosol cytoxicity assays was determined according to the equation: (*Np* × *Vp*)/*Vm* where *Np* is the number of puffs, *Vp* is the volume of 1 puff, and *Vm* is the volume of the medium.

### Identification and quantification of flavor chemicals in JUUL EC Pod Fluids and Aerosols

The pre-filled pod fluid obtained prior to aerosolization of the JUUL pod is referred to as “unvaped fluid”. The fluid left in the pod after the aerosol has been collected is referred to as “vaped fluid”. Unvaped fluids, vaped fluids and aerosols were analyzed using GC/MS. For each sample, 50 μL were dissolved in 0.95 mL of isopropyl alcohol (IPA) (Fisher Scientific, Fair Lawn, NJ). All diluted samples were shipped overnight on ice to Portland State University and analyzed using GC/MS on the day they were received. A 20 μL aliquot of internal standard solution (2000 ng/μL of 1,2,3-trichlorobenzene dissolved in IPA) was added to each diluted sample before analysis. Using internal standard-based calibration procedures described elsewhere,^22^ analyses for 178 flavor-related target analytes were performed with an Agilent 5975C GC/MS system (Santa Clara, CA). A Restek Rxi-624Sil MS column (Bellefonte, PA) was used (30 m long, 0.25 mm id, and 1.4 μm film thickness. A 1.0 μL aliquot of diluted sample was injected into the GC with a 10:1 split. The injector temperature was 235 °C. The GC temperature program for analyses was: 40 °C hold for 2 min; 10 °C/min to 100 °C; then 12 °C/min to 280 °C and hold for 8 min at 280 °C, then 10 °C/min to 230 °C. The MS was operated in electron impact ionization mode at 70 eV in positive ion mode. The ion source temperature was 220 °C and the quadrapole temperature was 150 °C. The scan range was 34 to 400 amu. Each of the 178 target analytes was quantitated using authentic standard material and an internal standard (1,2,3-trichlorobenzene) normalized multipoint calibration.

### Cell Culture

Human bronchial epithelial cells (BEAS-2B) obtained from the American Type Culture Collection (ATCC) were cultured in Airway Epithelial Cell Basal Medium (Manassas, VA) supplemented with 1.25 mL of human serum albumin, linoleic acid and lecithin (HLL supplement), 15 mL of L-glutamine, 2 mL of extract P, and 5.0 mL airway epithelial cell supplement (Manassas, VA). Nunc T-25 tissue culture flasks were coated overnight with basal medium, collagen, bovine serum albumin and fibronectin prior to culturing and passaging cells. At 90% confluency, cells were harvested using Dulbecco’s phosphate buffered saline (DPBS) for washing and incubated with 2 mL of 0.25% trypsin EDTA/DPBS and poly-vinyl-pyrrolidone for 3 mins at 37°C to allow detachment. Cells were cultured in T-25 flasks at 75,000 cells/flask, and the medium was replaced every other day. For the in vitro assays, cells were plated at 8,000 – 10,000 cells/well in pre-coated 96-well plates and allowed to attach overnight prior to a 24-hour treatment.

### Cell Viability and Cytotoxicity Assays

The toxicities of unvaped and vaped pod fluids and their resulting aerosol fluids were determined using three assays. Treatments were performed over 3-fold dilutions with the highest concentration being 10% for the fluids and 6 TPE solutions for the aerosols, which ranged from 1.3 to 3%. Serial dilutions in culture medium were arranged in 96-well plates with negative controls placed next to the highest and lowest concentration to check for a vapor effect.^18^ Cells were exposed for 24 hours before performing the MTT 3-(4,5-dimethylthiazol-2-yl)-2,5-diphenyltetrazolium bromide (MTT), Neutral Red Uptake, (NRU) and Lactate Dehydrogenase (LDH) assays.

The MTT cytotoxicity assay measures mitochondrial reductases which convert the water soluble MTT salt to a formazan that accumulates in healthy cells. Post 24-hours of treatment, 20 μL of MTT (Sigma-Aldrich, St Louis, MO) dissolved in 5 mg/mL of DPBS (Fisher Scientific, Chino, CA) were added to each well and incubated for 2 hrs at 37°C. Solutions were removed, and 100 μL of dimethyl sulfoxide (DMSO) (Fisher, Chino, CA) were added to each well and gently mixed on a shaker. The absorbance of control and treated wells was read against a DMSO blank at 570 nm using an Epoch micro-plate reader (Biotek, Winooski, VT). Each chemical was tested in three independent experiments.

The NRU assay measures the uptake of neutral red dye, which accumulates within the lysosomes of healthy living cells. A working solution of 4 μg of neutral red stock (4 mg NR/mL of PBS without Ca^2^+ and Mg^2^+) per mL of cell culture medium was prepared and incubated at 37°C overnight to dissolve the neutral red. Following exposure of cells to treatments, all medium was removed, and cells were incubated with 150 μL of neutral red solution for 2 hours. Cells were washed with PBS and 150 μL of lysis buffer (50% EtOH/ 49% deionized H_2_O/ 1% acetic acid) were added to each well and gently mixed to achieve complete dissolution. The absorbance of control and treated wells at 540 nm was recorded using an Epoch micro-plate reader (Biotek, Winooski, VT).

The LDH leakage assay measures the activity of lactate dehydrogenase released into the culture medium and is an indicator of cell death or cytotoxicity due to plasma membrane damage. Reagents and solutions were prepared using an in-house recipe developed by OPS Diagnostics (Sigma-Aldrich, St Louis, MO). 200 mM TRIS, pH 8 (22.2 g Tris-HCl and10.6 g Tris-base and 50 mM of lithium lactate (19.6 mg/mL) were prepared in water. Tetrazolium salt (INT) was dissolved in DMSO (33 mg/mL), phenazine methosulphate (PMS) was dissolved in water (9 mg/mL), and β-nicotinamide adenine dinucleotide (NAD) sodium salt was dissolved in water (3.7 mg/mL). All three reagents (INT, PMS and NAD) were used to make the INT/PMS/NAD solution. 50 μL of all reagents were added to 96-well plates followed by 50 μL of culture medium obtained from both treated and control. The absorbance of all wells was measured at 490 nm using an Epoch micro-plate reader (Biotek, Winooski, VT).

### Statistical Analyses

All cytotoxicity assays were carried out using three independent experiments, and each experiment had triplicate points. Data were statistically analyzed with a one-way analysis of variance (ANOVA), and each concentration was compared to the untreated control with Dunnett’s post hoc test using Prism software (GraphPad, San Diego). For the nicotine concentration data, means were analyzed using an ANOVA followed by Bonferroni’s post hoc test.

## RESULTS

### Identification of Flavor Chemicals in JUUL Pods

Fifty-nine of 177 flavor chemicals on our target list were identified and quantified in duplicates of the eight JUUL flavor pods (Figure 1). The duplicate data were generated using fluids from two different unvaped pods analyzed at different times. The total concentration of flavor chemicals in each product appears above each column. Abbreviations of JUUL pod names are on the *x*-axis, and safety classifications based on existing oral rat LD_50_ data^23^ are on the *y*-axis. Within each safety classification, the chemicals are ranked from the most to least potent. Rat oral toxicity data were used for ranking because they were available for most chemicals in the heat map, while inhalation LD_50_ data were seldom available for rats or humans. Forty-three of the 59 chemicals had concentrations >0.01 mg/mL, 13 were >0.1mg/ mL, and 3 (menthol, vanillin and ethyl maltol) were >1.0 mg/ mL. The highest concentrations of menthol, vanillin and ethyl maltol in unvaped pod fluids were 15, 6.9 and 1.8 mg/ mL, respectively. Duplicate pods were generally similar to each other, however, “Fruit Medley-1” contained five times the total flavor chemical concentration as its duplicate pod. The “Fruit Medley” sample with 0.3 mg/mL was similar to levels in the “Classic Tobacco” and “Virginia Tobacco” pods, which were all lower than 0.5 mg/mL.

**Figure 1.**
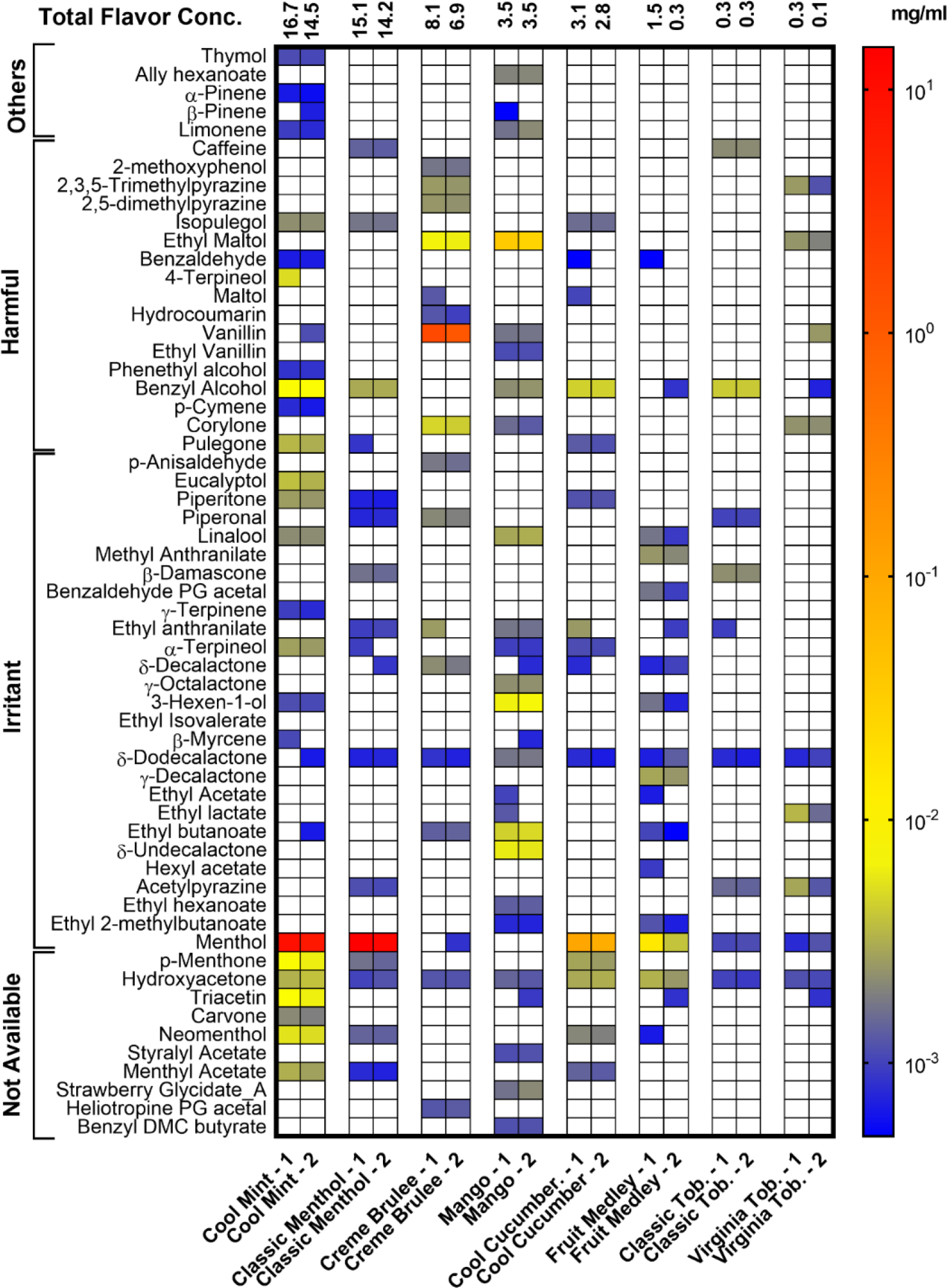
Heat map of flavor chemicals in eight duplicate JUUL pod fluids. Chemicals are ordered on the *y*-axis according to their toxicity (Others, Harmful, Irritant) based on LC50 data from rat oral exposures, and within each class, they are ranked from most to least toxic. The “Others” category on the *y*-axis represents chemicals that are corrosive, toxic, harmful, irritants as well as dangerous to the environment. JUUL products (*x*-axis) are ordered according to the total weight (mg/mL) of the flavor chemicals in each product with the highest concentration at the left. The total flavor chemical concentration (mg/mL) is indicated at the top of each column. The color gradient on the right shows the concentrations of the flavor chemicals in the heat map. Three chemicals (vanillin, ethyl maltol, and menthol) in the orange to red color gradient were ≥1 mg/mL in at least one product. JUUL pod code: Classic Tob. = “Classic Tobacco”; Virginia Tob. = “Virginia Tobacco”. The numbers 1 and 2 with the JUUL pod codes designate the first and second pod tested.

### Nicotine and Total Flavor Chemical Concentrations in EC Products

JUUL pods contain solvents, flavor chemicals, and varying concentrations of nicotine. The nicotine concentrations were evaluated in 183 EC products, which included the popular Vuse and JUUL products (Figure 2a). Nicotine concentrations in the EC fluids fell into one of three groups: (1) most products had 1.6 − 34.4 mg/mL (blue dots), (2) Vuse products had 18.9 − 38.8 mg/mL (green dots), and (3) JUUL had 59.2 − 66.7 mg/mL (red dots) (Figure 2a). The average concentration of nicotine was significantly higher in JUUL than in the other two groups (Figure 2b).

**Figure 2.**
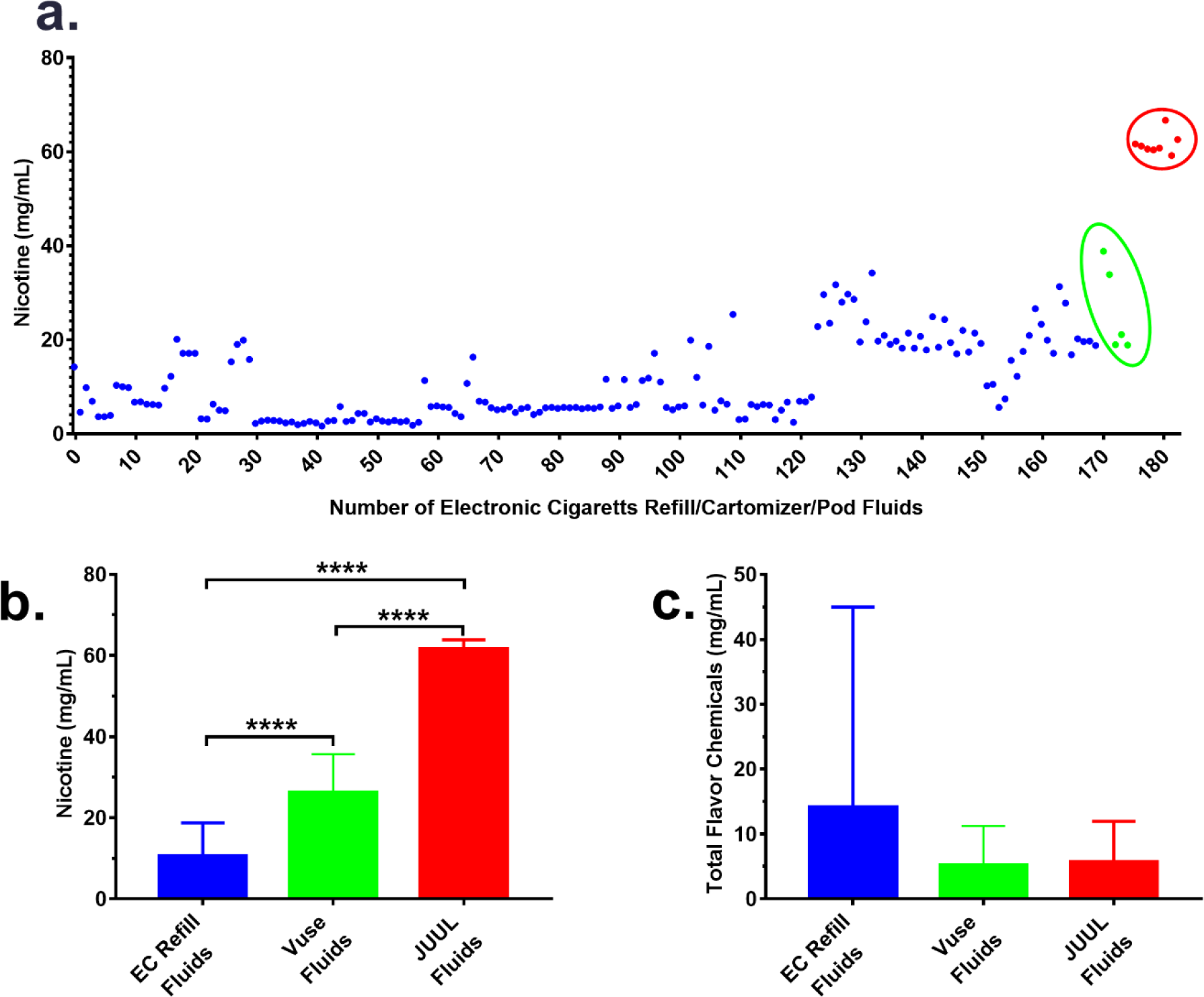
Nicotine and total flavor chemical concentrations in EC products. (a) Nicotine concentrations in 183 EC products. Red dots represent eight JUUL products; green dots represent 5 Vuse cartomizer fluids, and blue dots represent 170 refill fluids from 34 brands. The *y*-axis shows nicotine concentrations in each EC product listed on the *x*-axis. (b) The mean concentrations of nicotine in 170 EC refill fluids from 34 brands (blue bar), five Vuse cartomizers (green bar), and eight JUUL pods (red bar). The mean concentrations of nicotine were significantly different in each group. **** = p < 0.0001. (c) The mean concentrations of total flavor chemicals in 170 EC refill fluids from 34 brands (blue bar), five Vuse cartomizers (green bar), and eight pod JUUL pods (red bar).

The total concentration of flavor chemicals was compared in 183 EC products (170 refill fluids, five Vuse cartomizer fluids and eight JUUL pod fluids) (Figure 2c). Concentrations in refill fluids were highly variable and ranged from 0.1 to 362.3 mg/mL. In contrast, concentrations in cartomizers and pods were similar and generally lower than in refill fluids. Vuse cartomizers had total flavor chemical concentrations ranging from 0.7 to 15.7 mg/mL, while JUUL pods ranged from 0.2 to 15.6 mg/mL.

### Concentrations of Total Flavor Chemicals and Nicotine in JUUL Fluids and Aerosols

The total concentration of flavor chemicals in unvaped pod fluids, vaped fluids, and aerosols ranged between 0.1 – 16.7, 0.1 – 14.7, and 0.1 – 9.1 mg/mL, respectively (Figure 3a). Transfer from the fluid to the aerosol was variable, but in general was over 50% efficient. Only fluids from “Cool Mint” and “Classic Menthol” pods had total flavor chemical concentrations >10 mg/mL. “Crème Brulee”, “Mango”, “Cool Cucumber” and “Fruit Medley” had total flavor chemical concentrations between 0.3 and 8.1 mg/mL, while the two tobacco flavors had negligible concentrations.

**Figure 3.**
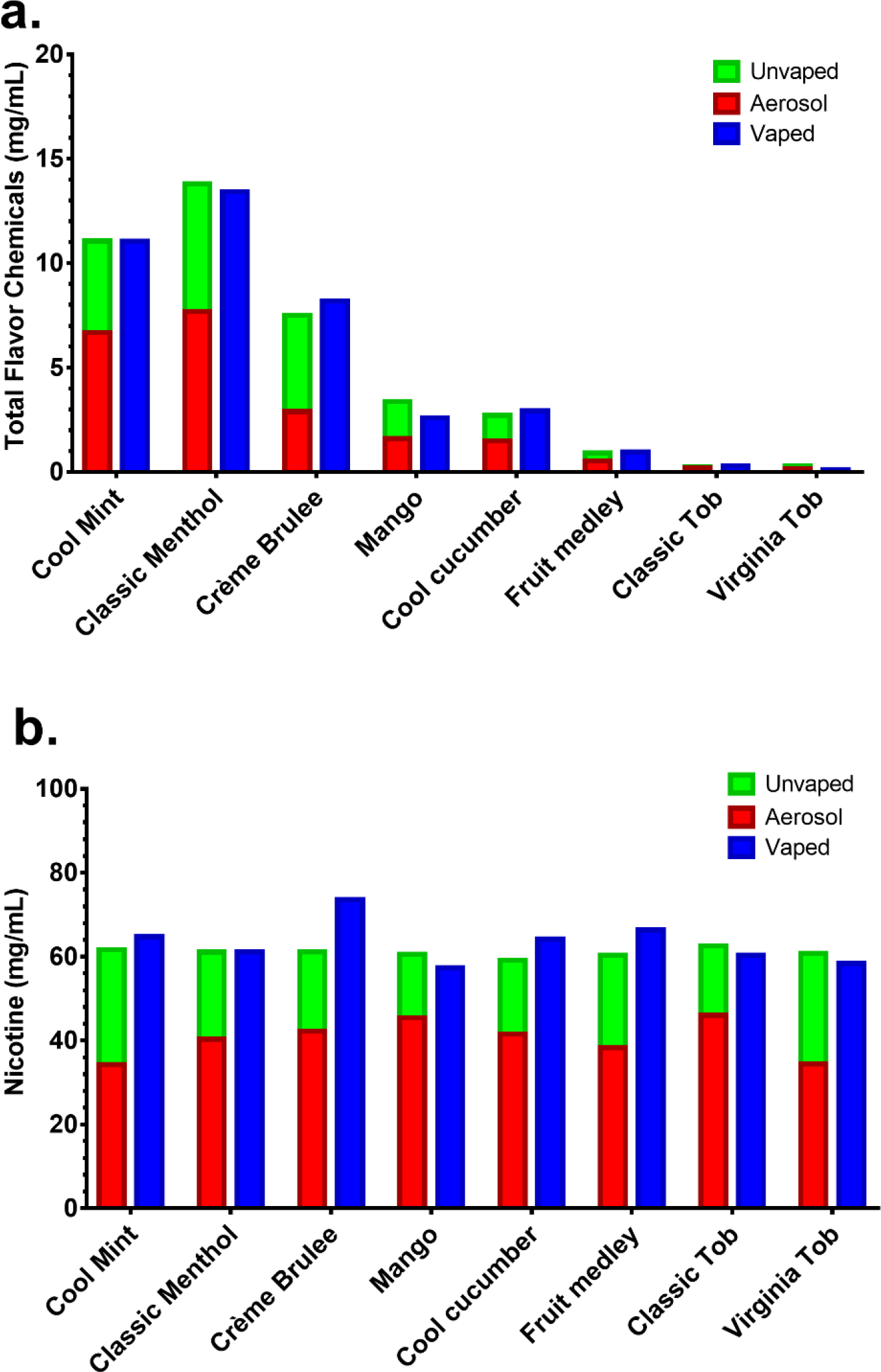
Total flavor chemical and nicotine concentrations in JUUL pod fluids and aerosols. (a) The total flavor chemical concentrations in unvaped pod fluids, vaped pod fluids, and aerosols. (b) Concentrations of nicotine in unvaped pod fluids, vaped pod fluids, and aerosols. The total flavor chemical concentrations and nicotine concentrations were very similar in the unvaped and vaped pod fluids. Each bar is mean concentration of two independent experiments.

In JUUL products, nicotine concentrations averaged 60.9 mg/mL, 63.5 mg/mL and 41.2 mg/mL in unvaped, vaped, and aerosol samples, respectively (Figure 3b). Transfer efficiently for nicotine to the aerosol was between 56 − 75%.

### Individual Flavor Chemicals and Transfer Efficiency

In comparison with other EC refill fluids that we have analyzed,^17^ JUUL uses a small number of different flavor chemicals in their pods (Figure 4). Five of eight products had 1-2 flavor chemicals (menthol, vanillin or ethyl maltol) >1 mg/mL, and these were generally present in about equal concentrations in both unvaped and vaped fluids. Menthol was the major flavor chemical in four of the flavor pods (“Cool Mint”, “Classic Menthol”, “Cool Cucumber” and “Fruit Medley”), although its concentration varied with “Classic Menthol” having the highest concentration (14.9 mg/mL) and “Fruit Medley” the lowest (0.7 mg/mL). Vanillin and ethyl maltol were the major flavor chemicals in “Crème Brulee” and “Mango”, respectively. “Classic Tobacco” had low levels of benzyl alcohol, while flavor chemicals were negligible in “Virginia Tobacco”. These major flavor chemicals in each product generally transferred well to the aerosol with transfer efficiencies ranging from 39 to 62%.

**Figure 4.**
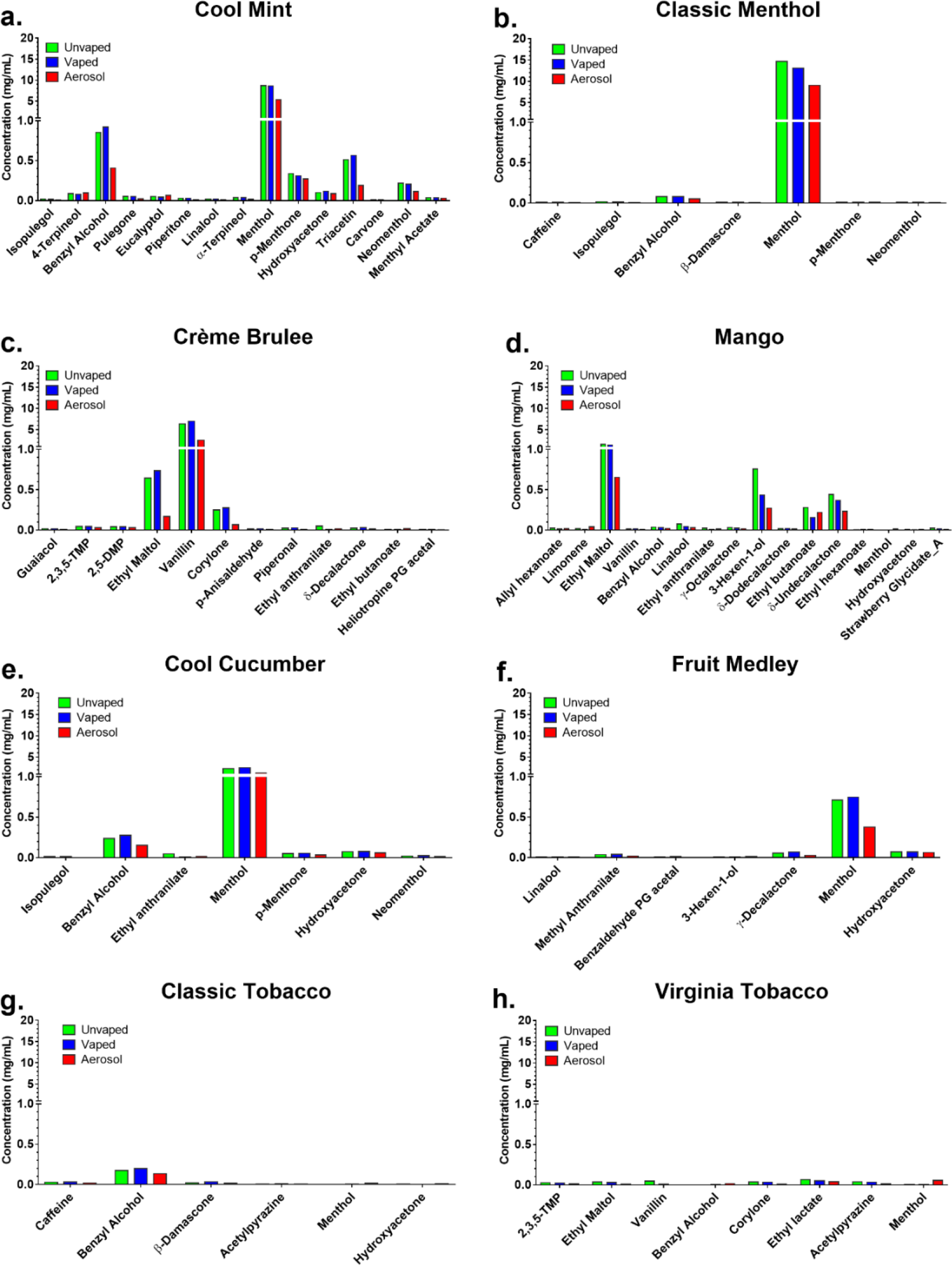
Concentrations of individual flavor chemicals in JUUL pod fluids and aerosols. (a) “Cool Mint”, (b) “Classic Menthol”, (c) “Crème Brulee”, (d) “Mango”, (f) “Fruit Medley”, (g) “Classic Tobacco”, and (h) “Virginia Tobacco”. Most fluids contained 1-2 flavor chemicals >1 mg/mL, except the tobacco flavored products, which had very low concentrations of flavor chemicals. Flavor chemicals >1mg/mL transferred from unvaped pod fluids into the aerosols with 39 to 62% efficiency. Each bar is the mean concentration of two independent experiments.

### Cytotoxicity of JUUL Pod Fluids and Aerosols

Cytotoxicities of both fluids and aerosols were evaluated with BEAS-2B cells using the MTT, NRU, and LDH assays. Products were considered cytotoxic if they produced an effect that was 30 % less than the untreated control (referred to as the IC_70_) in accordance with ISO protocol # 10993-5:2009(E) international standard.^24^ JUUL pod fluids were cytotoxic in both the MTT and NRU assays for all pod flavors. Generally, IC_70_s and IC_50_s were reached at fluid concentrations between 1-10%, and all products produced a maximum effect at 10% (Figures 5a-b and 5d-e). Cytotoxicity was also observed in the MTT and NRU assays when cells were tested with JUUL pod aerosols (Figures 5c and 5f). The highest aerosol concentration of 6TPE, when converted to percentage concentration of pod fluid, ranged from 1.3% to 3.0%. In the MTT assay, IC_70_s for aerosols varied with different pod flavors and generally were reached between concentrations of 0.31% to a 1.8% (Table 1). All aerosols were cytotoxic in the MTT assay according to the ISO standard and produced an IC_70_ at the high dose. In the NRU assay, IC_70_s were reached for five of the eight JUUL flavor pods (Table 2). Aerosols from three flavors pods (“Classic Menthol”, “Classic Tobacco”, and “Virginia Tobacco”) did not produce a significant effect.

**Figure 5.**
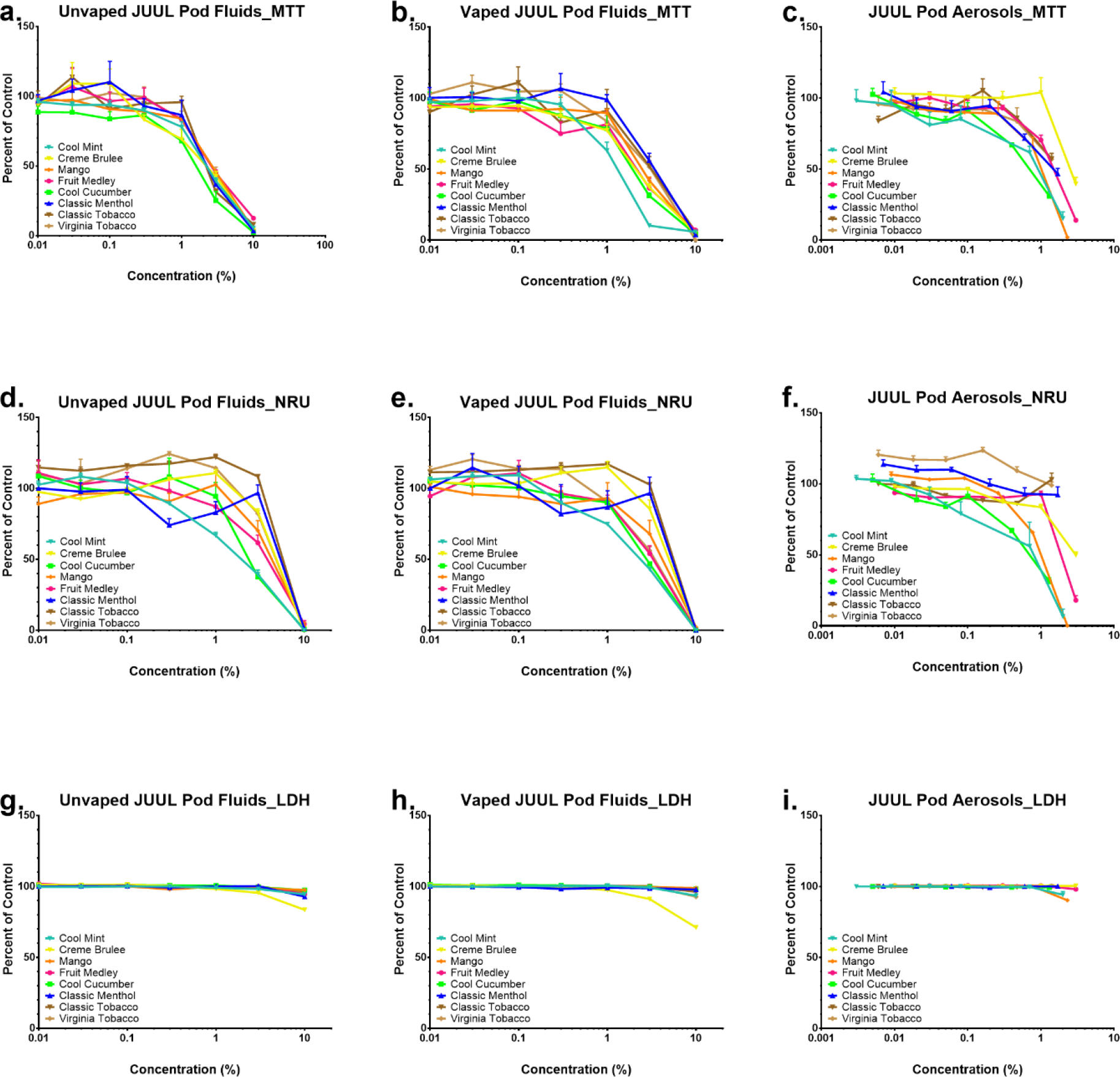
Concentration-response curves for BEAS-2B cells treated with JUUL pod fluids and aerosols. (a-c) MTT assay, (d-f) NRU assay, and (g-i) LDH assay for all eight pod variants. The *y*-axis shows the response of cells in each assay as a percentage of the untreated control. Each point is the mean ± standard error of the mean for three independent experiments.

**Table 1.** IC_70_ and IC_50_ (mg/mL) of JUUL pod fluids and aerosols in the MTT Assay.

| JUUL Pod Flavors<sup>a</sup> | Unvaped Fluids |  | Vaped Fluids |  | Aerosols |  |  |
| --- | --- | --- | --- | --- | --- | --- | --- |
|  | IC<sub>70</sub> | IC<sub>50</sub> | IC<sub>70</sub> | IC<sub>50</sub> | IC<sub>70</sub> | IC<sub>50</sub> | Highest Conc. (%) |
| “Cool Mint” | 0.92 | 2.17 | 0.79 | 1.25 | 0.31 | 0.64 | 2 |
| “Cool Cucumber” | 1.10 | 1.43 | 1.23 | 1.93 | 0.33 | 0.68 | 1.3 |
| “Mango” | 1.52 | 2.57 | 1.61 | 2.58 | 0.65 | 0.93 | 2.3 |
| “Classic Menthol” | 1.48 | 2.33 | 2.14 | 3.24 | 0.67 | 1.51 | 1.7 |
| “Virginia Tobacco” | 1.54 | 2.61 | 1.66 | 2.88 | 0.85 | 2.17 | 1.4 |
| “Classic Tobacco” | 1.60 | 2.37 | 1.87 | 3.07 | 0.89 | 1.67 | 1.4 |
| “Fruit Medley” | 1.52 | 2.70 | 1.35 | 2.00 | 1.01 | 1.42 | 3 |
| “Creme Brulee” | 1.03 | 1.97 | 1.27 | 2.06 | 1.80 | 2.90 | 3 |

<sup>a</sup> = order of pod flavors ranked according to IC<sub>70</sub> of aerosols

**Table 2.** IC_70_ and IC_50_ (mg/mL) of JUUL pod fluids and aerosols in the NRU Assay.

| JUUL Pod Flavors<sup>a</sup> | Unvaped Fluids |  | Vaped Fluids |  | Aerosols |  |  |
| --- | --- | --- | --- | --- | --- | --- | --- |
|  | IC<sub>70</sub> | IC<sub>50</sub> | IC<sub>70</sub> | IC<sub>50</sub> | IC<sub>70</sub> | IC<sub>50</sub> | Highest Conc. (%) |
| “Cool Mint” | 1.32 | 1.81 | 1.21 | 2.18 | 0.20 | 0.54 | 2 |
| “Cool Cucumber” | 1.65 | 2.55 | 1.70 | 2.77 | 0.42 | 0.68 | 1.3 |
| “Mango” | 3.08 | 3.75 | 1.61 | 3.75 | 0.65 | 0.89 | 2.3 |
| “Fruit Medley” | 2.29 | 3.50 | 1.35 | 3.11 | 1.39 | 1.98 | 3 |
| “Creme Brulee” | 3.68 | 4.88 | 3.75 | 5.07 | 1.52 | 3.23 | 3 |
| “Classic Menthol” | 4.28 | 5.09 | 2.14 | 4.30 | > 1.7 | > 1.7 | 1.7 |
| “Classic Tobacco” | 4.82 | 7.94 | 1.87 | 7.84 | > 1.4 | n/a | 1.4 |
| “Virginia Tobacco” | 3.69 | 4.91 | 1.66 | 3.21 | n/a | n/a | 1.4 |

<sup>a</sup> = order of pod flavors ranked according to IC<sub>70</sub> of aerosols

With JUUL pod fluids and aerosols, little effect was seen in the LDH assay (Figures 5g-i), indicating that in general, fluids and aerosol treatments did not cause rupture of BEAS-2B plasma membranes.

### Correlation between Nicotine Concentration, Flavor Chemical Concentration, and Toxicity

Since flavor chemicals can cause cytotoxicity, especially at concentration >1 mg/mL,^17^ linear regression analyses were performed to parse out the relative contribution of nicotine, total flavor chemicals, and individual flavor chemicals to the cytotoxicity observed with JUUL pod fluids and aerosols (Figures 6 and 7). For unvaped JUUL fluid, there was a high correlation between cytoxicity (percent of untreated control) and the concentration of nicotine plus total flavor chemicals in both the MTT (R^2^ = 0.871; p<0.0001) and NRU (R^2^ = 0.861; p<0.0001) assays (Figure 6a). When nicotine and flavor chemical concentrations were analyzed separately (Figures 6b and 6c), the correlation coefficient for nicotine concentrations alone versus cytotoxicity (R^2^ = 0.879 for MTT) was almost equivalent to that of nicotine and flavor chemicals concentrations combined (R^2^ = 0.871 for MTT). In contrast, total flavor chemical concentration alone (without nicotine) was only moderately/weakly correlated to cytoxicity (R^2^ = 0.379 for MTT and 0.383 for NRU), nevertheless the correlation was significant (p<0.0001 for both MTT and NRU). The correlation between cytotoxicity and the concentrations of individual flavor chemicals found at concentrations >1 mg/mL was moderate for ethyl maltol and weak for menthol and vanillin (Figures 6d-f); nevertheless, all correlations were statistically significant (Figures 6d-f). A similar pattern of linear correlation and statistical significance was observed with vaped fluids in both the MTT and neutral assays (Supporting Information, S2).

**Figure 6.**
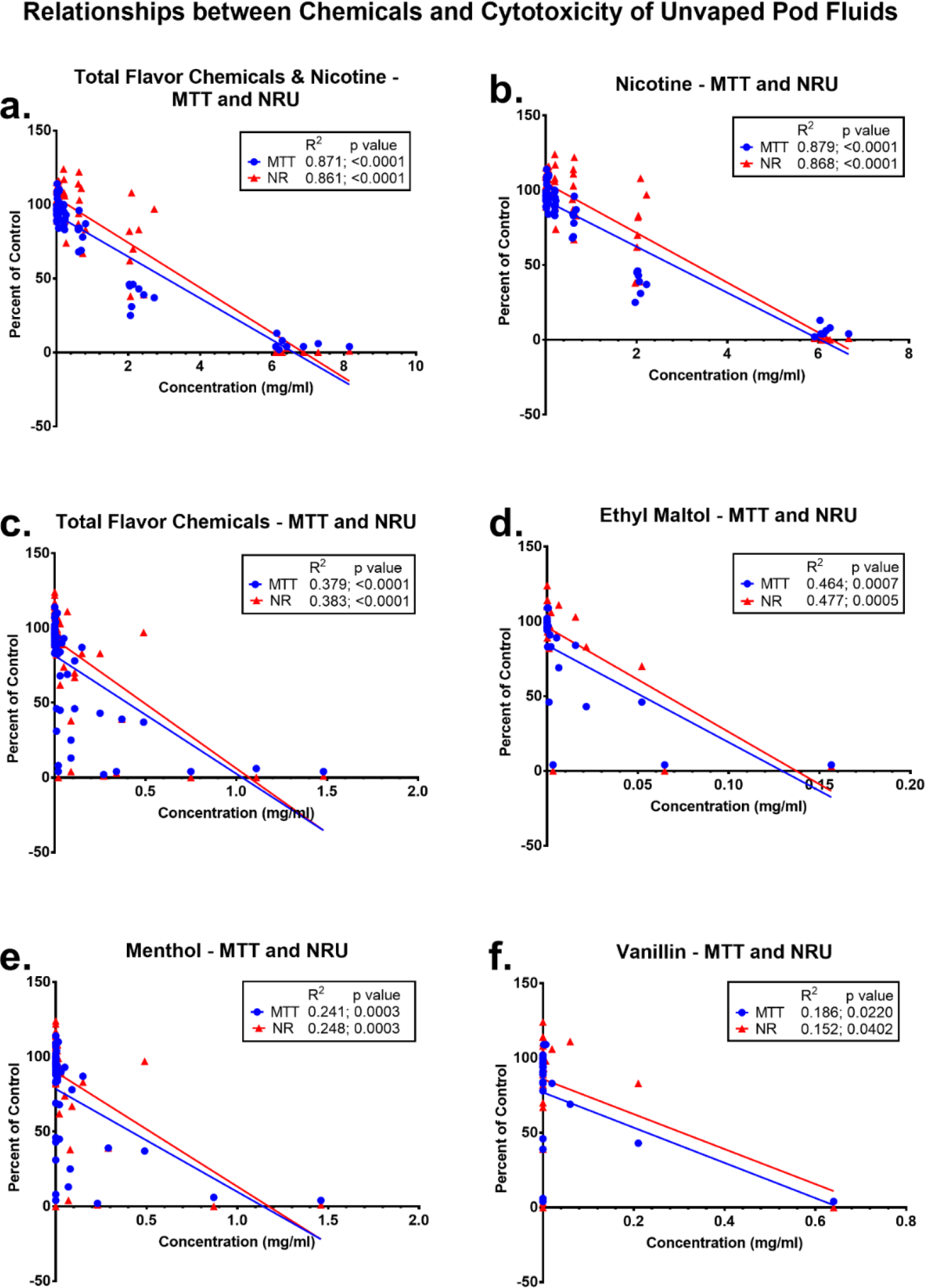
Relationship between cytoxicity of unvaped pod fluids and concentrations of nicotine and the flavor chemicals. Linear regression analysis for cytotoxicity (*y*-axis, expressed as a percentage of the untreated control) in the MTT and NRU assays versus the concentrations of: (a) total flavor chemicals and nicotine, (b) nicotine only, (c) total flavor chemicals only, (d) ethyl maltol, (e) menthol, and (f) vanillin. Blue dots and red triangles represent concentrations tested in the MTT and NRU assay, respectively. Cytotoxicity was strongly correlated with total concentration of chemicals (flavor chemicals and nicotine) and with nicotine concentration only and weakly to moderately correlated with the concentrations of total flavor chemicals, ethyl maltol, menthol and vanillin. All correlations were significant (p<0.05).

For JUUL aerosols, correlations between cytotoxicity and total chemicals (nicotine plus flavor chemicals) (Figure 7a), nicotine alone (Figure 7b), and ethyl maltol (Figure 7d) were strong (R^2^ > 0.75, except for two NRU R^2^s which were > 0.45) and significant (all p<0.0001) (Figures 7a-b, and 7d). Flavor chemicals alone (Figure 7c) and menthol (Figure 7e) were weakly correlated to cytotoxicity (R^2^ ranged from 0.099 to 0.361), while R^2^ for vanillin was weak and not significant (p>0.05) (Figure 7f).

**Figure 7.**
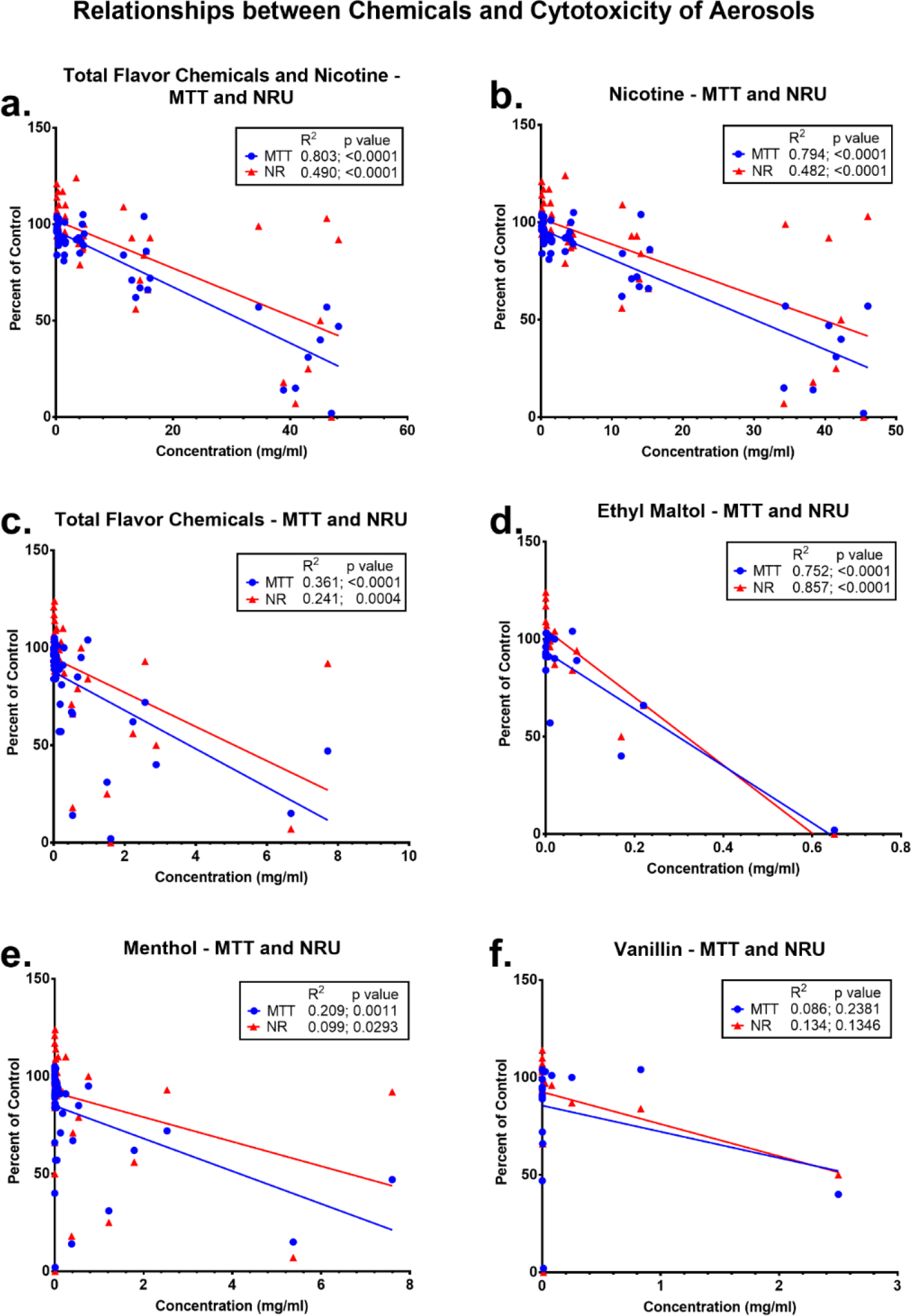
Relationship between cytoxicity of pod aerosols and the concentrations of nicotine and the flavor chemicals. Linear regression analysis for cytotoxicity in the MTT and NRU assays versus the concentrations of: (a) total flavor chemicals and nicotine, (b) nicotine only, (c) total flavor chemicals only, (d) ethyl maltol, (e) menthol, and (f) vanillin. Blue dots and red triangles represent the concentrations tested in the MTT and NRU assay. Cytotoxicity (percent of control) was strongly correlated with the total concentration of chemicals (flavor chemicals and nicotine), nicotine concentration only, and ethyl maltol concentration. The correlations between cytotoxicity and the concentrations of total flavor chemicals and menthol were moderate and weak, respectively. The correlation between cytotoxicity and vanillin concentration was not significant.

## DISCUSSION

While the health complications associated with EC use are appearing in case reports and the infodemiological literature,^25,26^ to date no health reports have been made for consumers of JUUL products. Nicotine concentrations were higher in JUUL pod fluids than in any of the 183 EC refill and cartomizer fluids that we have examined previously^27^ (Figure 2a). Concentration-response curves for the JUUL fluids were remarkably similar among the flavor pods and reached a maximum effect in the MTT and NRU assays at a 10% concentration for all samples. Aerosols were all cytotoxic at concentrations >1 %. Cytotoxicity of aerosols was strongly correlated with total chemical concentrations, nicotine concentration, and ethyl maltol concentration, which was 1.81 mg/mL in one JUUL product. While we have previously reported that the concentrations of some flavor chemicals in some EC products are high enough to be cytotoxic,^19,20^ JUUL pods are the only EC product that we have studied in which cytotoxicity can be attributed to the concentrations of both nicotine and a flavor chemical (ethyl maltol).

Only 1-2 flavor chemicals were present at concentrations >1 mg/mL in each JUUL product, similar to some refill fluids from other manufacturers that contained 1-4 flavor chemicals/product at 1 mg/mL or greater.^17^ In general, the concentrations of individual flavor chemicals in JUUL products were relatively low compared to other cartomizer style EC and refill fluids.^16,17^ Two exceptions were JUUL “Cool Mint” and “Classic Menthol”, which both had menthol concentrations >10 mg/mL. Others have reported that the minty flavors may be the most popular of the JUUL products,^3^ which could be due to a stronger flavor imparted by their high concentrations of menthol or the effects of menthol on nicotine metabolism.^28^ In contrast to the minty products, the two JUUL tobacco-flavored pods had very low concentrations of flavor chemicals. It is possible that the high concentration of nicotine and acid in JUUL pods imparts some flavor features to the aerosol making the use of additional chemicals unnecessary in the “Classic Tobacco” and “Virginia Tobacco” pods or that the predominant aroma molecules for those flavor profiles were not included in the GC/MS target compounds. The low levels of flavor chemicals in most JUUL pods may reduce their odor, which would facilitate “stealth” use, a desirable feature among middle and high school students who vape in class or in rest rooms.^12^

The flavor chemicals that were present in JUUL pods at very low concentrations are likely co-constituents of the major flavor chemicals (i.e., menthol, vanillin and ethyl maltol) or may in some cases be added to impart subtle flavor accents. With respect to manufacturing practices, duplicate pods and packages were identical and contained similar flavor chemicals. However, during aerosol production, pods did not perform uniformly on the smoking machine, and some pods did not work at all. This inconsistency in puff production may also account for the relatively low transfer efficiencies seen with some pods.

Nicotine concentrations in the JUUL products were significantly higher than in any other EC cartomizers and refill fluids our laboratory has evaluated (total 183).^27,29^ The average nicotine concentration in JUUL pods in our study (60.9 mg/mL) agrees well with an earlier report (61.6 mg/mL).^14^ A single JUUL pod contained more nicotine (56 - 66 mg) than a pack of cigarettes (2 mg/stick * 20 sticks = 40 mg/pack). The high concentrations of nicotine in JUUL EC is coupled to a high concentration of benzoic acid, which protonates nicotine making it less harsh when inhaled by users.^14,15^ The combination of the high nicotine concentration and its protonation by benzoic acid likely facilitates JUUL use and subsequent addiction, especially of adolescent or naïve consumers of JUUL products. The potential for addiction to JUUL products is compounded by the report that only 37% of the past 30-day consumers were aware that JUUL products always contain nicotine.^30^ In contrast to nicotine, total flavor chemical concentrations were not unusually high in JUUL pods and were found over a relatively narrow range of concentrations (15.7/mL being the highest). Refill fluids, in contrast, have a much wider range of total flavor chemical concentrations with the highest we have detected being 362.3 mg/mL. Moreover, the high concentrations of flavor chemicals are cytotoxic when tested in vitro.^17^ In this study, only one flavor chemical (ethyl maltol) was correlated with cytoxicity, as discussed below.

JUUL fluids and aerosols produced no significant effects in the LDH assay. Since this assay measures the release of LDH, a cytoplasmic enzyme, it is probable that treatment did not lyse cells or cause significant damage to the plasma membrane. In contrast, all pod fluids and most aerosols produced a cytotoxic response at a 10% concentration in the MTT and NRU assays. Our linear regression analysis showed that the nicotine and ethyl maltol concentrations in JUUL aerosols were high enough to account for most of the cytotoxicity observed with the MTT and NRU. Since nicotine concentrations were similar in all JUUL products and since cytoxicity can be attributed mainly to nicotine, the concentration-response curves for JUUL fluids were all similar. In some prior work with other EC products that had lower nicotine concentrations, cytotoxicity was correlated with the favor chemical concentration, not nicotine.^17,18,31^ Ethyl maltol concentration, which was also strongly correlated with aerosol cytotoxicity, was highest in the Mango pods (1.57 mg/mL), which were more potent than “Crème Brulee” and “Virginia Tobacco” (Figures 5c, and 5f), which both had lower concentrations of ethyl maltol (0.65 mg/mL and 0.03mg/mL, respectively) (Figure 1).

In the NRU assay, the “Classic Menthol” and “Classic Tobacco” aerosol did not inhibit uptake relative to the control. This could be because the concentrations of the aerosol did not reach 10%, as they did with fluids. In addition, these were the only flavors that contained caffeine (Figure 1), which is a stimulant. The caffeine concentrations in “Classic Menthol” and “Classic Tobacco” aerosols were 0.037mM and 0.090 mM, respectively. These concentrations are similar to those reported to provide protection to cells in other models^32^ and may explain our results with “Classic Menthol” and “Classic Tobacco” aerosol.

In summary, JUUL has previously raised two major concerns for the FDA. The first is the likelihood that JUUL use, which is widespread among middle school and high school students, will addict a new generation of adolescents to nicotine. The second it that these adolescents will eventually migrate to more dangerous tobacco products, such as conventional cigarettes. Our data clearly identify a third concern related to the high nicotine concentration in JUUL products, i.e., the potential for nicotine, as well as flavor chemicals such as ethyl maltol, to damage or even kill cells at the concentrations used in JUUL pods. Our exposures were acute and produced a maximal cytotoxic response that was strongly correlated with nicotine and ethyl maltol concentration. Maximal responses were observed at 10% concentrations of pod fluids, which would be lower than JUUL users inhale. It will be important in future work to determine if JUUL products, and other products containing nicotine salts, have adverse effects on consumers and if such effects lead to health problems with chronic use. In the meantime, the FDA could limit nicotine and flavor chemical concentrations in EC products

## Supporting information

## Author Contributions

PT, JFP and EO formed the concept and design of this study. Sample preparation, data collection and data processing were carried out by EO, WL, and KJM. Data were analyzed and interpreted by EO, WL, JFP, and PT. Writing and editing of the manuscript was done by EO, WL, KJM, JFP, and PT.

## Acknowledgment

We thank Riaz Golshan, Vivian To, Sara Leung, and Madeleine Vega for helping with aerosol production.

