## Supplementary material for "Toxicity of JUUL Fluids and Aerosols Correlates Strongly with Nicotine and Some Flavor Chemical Concentrations"

### Supporting Information

S1 Images of the eight JUUL Pod flavor evaluated in this study.

S2 Relationship between cytoxicity of vaped pod fluids and concentrations of nicotine and the flavor chemicals.

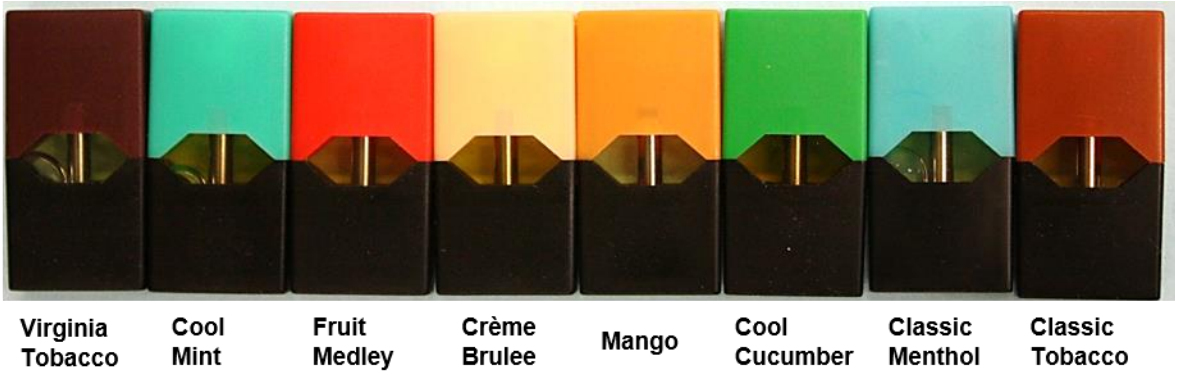


**Supplementary Figure 1.** Images of eight JUUL Pod Flavors evaluated in this study.

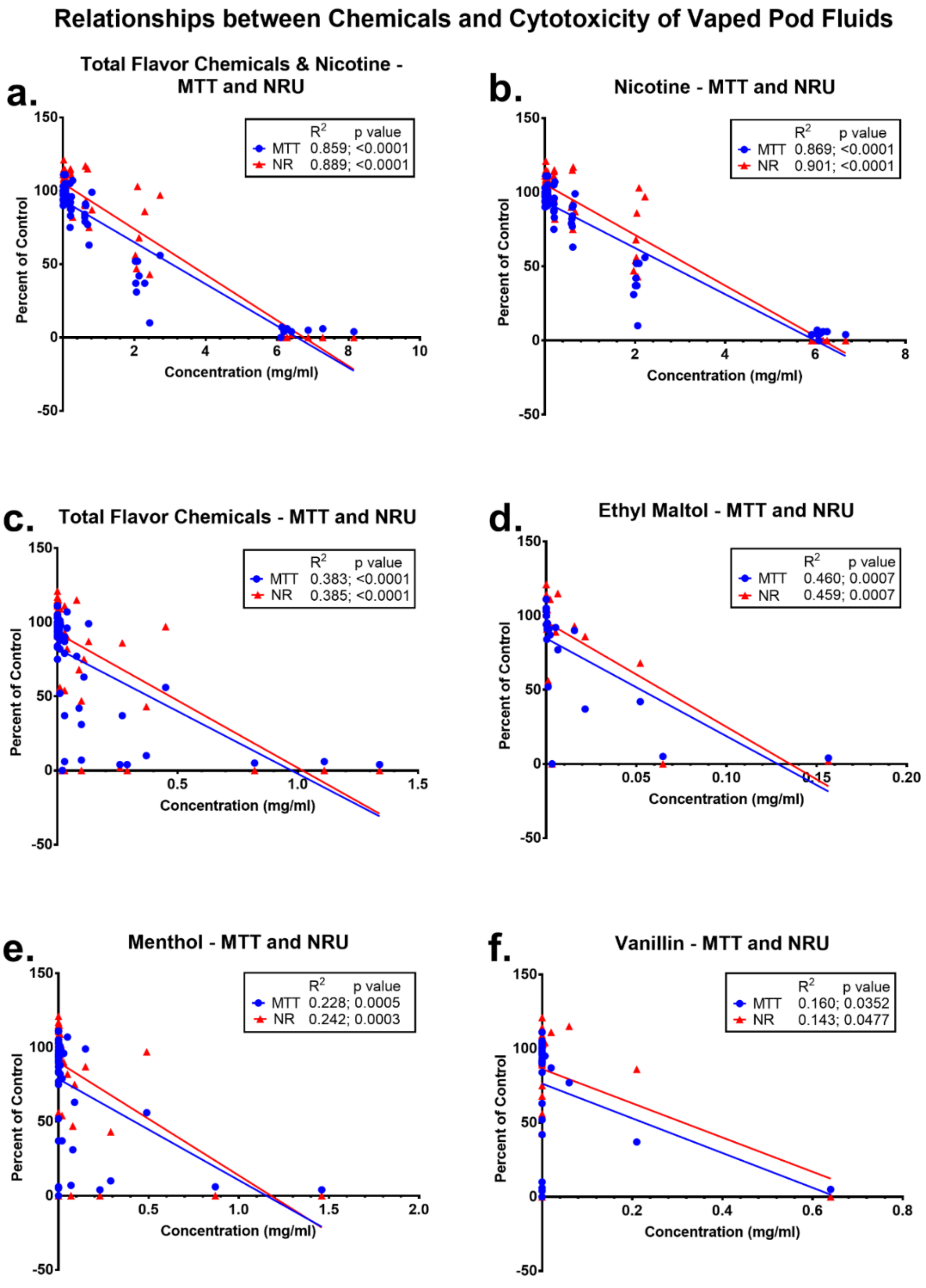


**Supplementary Figure 2.** Relationship between cytoxicity of vaped pod fluids and concentrations of nicotine and the flavor chemicals. Linear regression analysis for cytotoxicity (percentage of the untreated control) in the MTT and NRU assays versus the concentrations of: (a) total flavor chemical and nicotine, (b) nicotine only, (c) total flavor chemicals only, (d) ethyl maltol, (e) menthol, (f) vanillin. Blue dots and red triangles represent the concentrations tested in the MTT and NRU assay. Cytotoxicity was strongly correlated with the total concentration of chemicals (flavor chemicals and nicotine) and with nicotine concentration only and weakly to moderately correlated with the concentrations of total flavor chemicals, ethyl maltol, menthol, and vanillin. All correlations were significant ($p<0.05$).
